# Feature-based in-silico model to predict the *Mycobacterium tuberculosis* bedaquiline phenotype associated with Rv0678 variants

**DOI:** 10.64898/2026.02.12.705543

**Authors:** Wilma Quispe Rojas, Miguel de Diego Fuertes, Vincent Rennie, Emmanuel Riviere, Mahdi Safarpour, Annelies Van Rie

**Author notes:** Corresponding author: Annelies Van Rie.

## Abstract

Bedaquiline resistance is emerging globally and threatens the effectiveness of the novel short all-oral regimens for rifampicin-resistant tuberculosis. Following a systematic literature review, we quantified 13 sequence, biochemical, and structural features of 62 *Rv0678* missense variants reported in 136 *Mycobacterium tuberculosis* isolates. Using rigorous machine learning methods, we show that the strongest contributing features were the evolutionary conservation score and the shortest atomic distance to key functional sites. The final 5-feature model had good performance (ROC-AUC 0.826) and classified the bedaquiline phenotype with high accuracy [sensitivity 87.1% (95% CI, 78.3–92.6) and specificity 88.2% (95% CI, 76.6–94.5)]. Performance was lower in external validation, likely due to the measurement error introduced when using diverse phenotypic methods. missense variants on the mmpR5 protein structure and function. Integrating the five-feature *in-silico* in variant interpretation software could improve the prediction of the effect of *Rv0678* variants and guide clinical management of rifampicin-resistant tuberculosis.

## INTRODUCTION

The occurrence of more than 400,000 cases of rifampicin-resistant tuberculosis (RR-TB) annually hinders tuberculosis control efforts and burdens patients and health systems^1^. Bedaquiline is a core drug for the all-oral short regimens recommended by the World Health Organization (WHO) for treatment of RR-TB^2^.

Resistance to bedaquiline was reported soon after its introduction into routine care^3^, and is now considered ubiquitous^4^. The primary mechanism of bedaquiline resistance in *Mycobacterium tuberculosis (Mtb)* is the upregulation of the mmpS5/mmpL5 efflux system, which plays a crucial role in emitting bedaquiline from the mycobacterial cells. The mmpS5/mmpL5 operon is regulated by the transcriptional repressor mmpR5 protein, which is encoded by the *Rv0678* gene^5^. Overexpression of the efflux pump results in bedaquiline resistance through a reduced intracellular concentration of bedaquiline ^6^.

Many variants in the *Rv0678* gene have been identified in *Mtb* isolates, with variable impact on the bedaquiline phenotype. ^6^ This poses challenges to determining the statistical genotype-phenotype association for *Rv0678* variants. In the WHO catalogue of mutations in *Mtb* complex, most missense *Rv0678* variants are classified as “of uncertain significance” ^7^. This complicates the use of next-generation sequencing (NGS) for determining bedaquiline resistance in a clinical care context.

Feature-based machine learning models have been successfully used to assess the impact of genomic variants on structural protein and to predict resistance in *Mtb* to rifampicin, pyrazinamide, ethambutol, isoniazid and ofloxacin ^7–11^. For bedaquiline, three *in-silico* models have been published. One focused on missense variants in the *atpE* gene, which occur rarely in clinical samples ^12^. Another model focused on missense variants in *Rv0678*, but model accuracy was too low for clinical decision-making ^13^. A third model applied a convolutional neural network to all bedaquiline resistance-associated genes simultaneously, but its accuracy was also too low for clinical application ^14^.

In this study, we aimed to develop an accurate *in-silico* model to predict the effect of missense variants in *Rv0678* on the bedaquiline phenotype.

## MATERIAL AND METHODS

The development of the *in-silico* model for prediction of the bedaquiline phenotype followed a multi-step process (Fig. 1).

**Figure 1.**
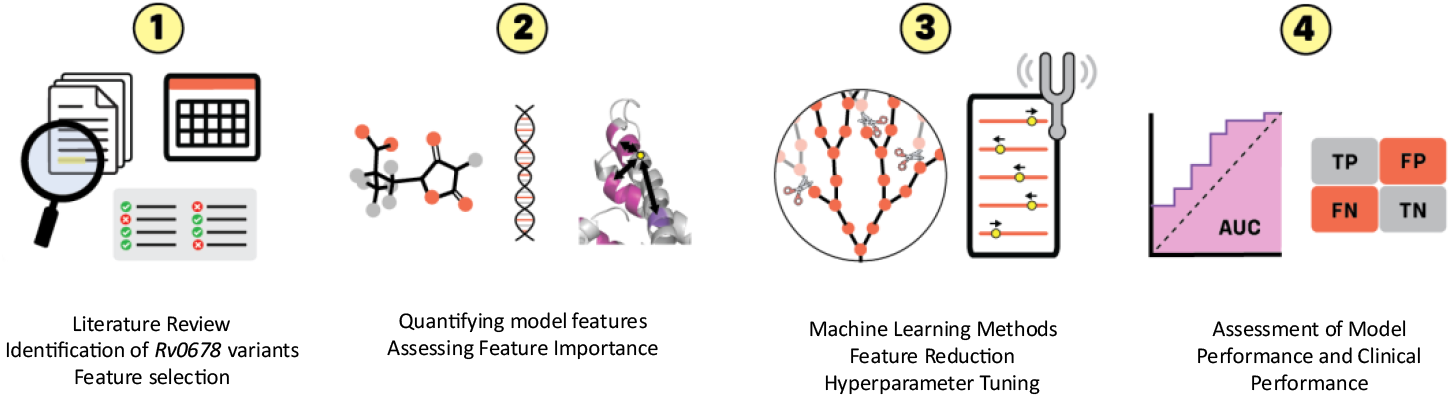
Multi-step process for the development of the *in-silico* model for prediction of the bedaquiline phenotype from genotypic data.

### Systematic literature review and data extraction

We performed a systematic literature review to identify all studies published between 1 January 2008 and 19 January 2026 that present paired genotypic-phenotypic data for bedaquiline in clinical *Mtb* isolates. We updated a prior literature review (1 Jan 2008 to December 2021) ^15^, using Rayyan web software ^16^, and applied the same search engines (PubMed Central, Europe PubMed and Scopus), search terms, methodology, and eligibility criteria.

For all isolates listed in eligible articles, we extracted the isolate study identification number, phenotypic drug susceptibility (DTS) result, minimum inhibitory concentration (MIC), variants present in the *atpE, Rv0678, pepQ, Rv1979, Rv0676, Rv0677* and *Rv1919c* genes, and the type of *Rv0678* variant detected.

### Selection of isolates and variants to create analytic datasets

For the main analysis, isolates were included if phenotypic testing was performed on the Mycobacteria Growth Indicator Tube (MGIT) platform using 1 μg/ml as the critical concentration ^17^ and the isolate contained a single missense variant in *Rv0678* without a co-occurring *atpE* variant. When information on *atpE* was missing, the isolate was assumed to be *atpE* wild-type, given the low prevalence of *atpE* variants in clinical isolates^18^. Isolates with a missense variant in *Rv0678* affecting amino acids (AA) in positions 1-14 or 160-165 were excluded as these positions are not present in the mmpR5 crystal structure^5^.

For each variant, we created a binary phenotype (susceptible or resistant) by assigning the ‘majority’ phenotype. Variants with an equal number of isolates reported as susceptible and resistant were excluded from the analysis.

### Identification of functional sites in the mmpR5 3D protein structure

To improve the prior *in-silico* model for *Rv0678* missense variants^13^, we added a feature of the distance to a functional site as was implemented in the *atpE* model^12^. We included the DxR motif (residues 88–92) and the alpha-helix recognition site (residues 62–68) as functional sites^19^. To identify additional candidate functional sites, we mapped the positions in the 3D DNA-bound mmpR5 conformation of all *Rv0678* variants occurring in phenotypically resistant *Mtb* isolates reported in ^19^ and used an iterative approach to identify 3D hotspots. To evaluate the biological relevance of the newly identified candidate functional sites, we assessed their proximity to functionally important regions of mmpR5 ^5^, including salt bridges, aromatic residues, and helix-capping motifs.

### Quantification of sequence, structural, and biochemical features to characterize the effect of missense variants in *Rv0678* on mmpR5 protein structure and function

To assess the effect of *Rv0678* variants on the 3D structure of the mmpR5 protein, we quantified 13 features of interest (Table 1) for each variant retrieved by our literature review. Analysis was performed in the ligand-bound conformation derived from the MmpR5 crystal structure (PDB 4NB5)^5^ and/or the DNA-bound conformation ^19^.

**Table 1:** Sequencing, structural, and biochemical features to characterize missense variants in *Rv0678*.

|  | Tool description | Value Range | Ref |
| --- | --- | --- | --- |
| **Sequencing features** |  |  |  |
| Evolutionary Conservation Score | ConSurf scores how conserved each AA is based on a multiple sequence alignment of homologous proteins | $\leq -1.0$ (highly conserved) to $\geq +1.5$ (highly variable) | 20 |
| Similarity between protein sequences | BLOSUM62 is a scoring matrix used to evaluate the likelihood of AA substitutions during evolution | $\sim -4$ (radical) to $+11$ (very conservative) | 21 |
| **Biochemical feature** |  |  |  |
| Chemical difference between AAs | Grantham distance quantifies biochemical differences between AAs based on 3 chemical properties: composition, polarity, and molecular volume | 0 (conservative) to $>150$ (radical) | 22 |
| **Structural features** |  |  |  |
| Relative Solvent Accessibility in DNA conformation | mCSM PPI assesses the relative solvent accessibility by measuring how exposed a residue is to solvent compared to its maximum possible exposure in an unfolded or reference state | 0 (Buried) to $>0.5 - 1$ (Exposed) | 23 |
| Relative Solvent Accessibility in ligand conformation |  |  |  |
| Protein Stability in DNA conformation | mCSM-Stability predicts how a missense mutation affects the thermodynamic stability of a protein | Positive $\Delta\Delta G$ is stabilizing; $\Delta\Delta G \approx 0$ minimal or no effect; negative $\Delta\Delta G$ is destabilizing | 23 |
| Protein Stability in ligand conformation |  |  |  |
| Protein-Protein interaction in DNA conformation | mCSM-PPI2 predicts the effect of point mutations on protein-protein interaction affinities | Positive $\Delta\Delta G$ is stabilizing; $\Delta\Delta G \approx 0$ minimal or no effect; negative $\Delta\Delta G$ is destabilizing | 24 |
| Protein-Protein interaction in ligand conformation |  |  |  |
| Protein dynamics in DNA conformation | DynaMut predicts the impact of point mutations on protein stability and dynamics by integrating structural bioinformatics with normal mode analysis | Positive $\Delta\Delta G$ is stabilizing; $\Delta\Delta G \approx 0$ minimal or no effect; negative $\Delta\Delta G$ is destabilizing | 25 |
| Protein dynamics in ligand conformation |  |  |  |
| Protein-DNA affinity in DNA conformation | The DNA_mCSM tool predicts how point mutations in proteins affect their binding affinity to DNA | Positive $\Delta\Delta G$ is stabilizing; $\Delta\Delta G \approx 0$ minimal or no effect; negative $\Delta\Delta G$ is destabilizing | 23 |
| Shortest Atomic Distance to one of five functional sites in DNA conformation | The shortest atomic distance from a mutation site to 5 key functional sites: alpha helix recognition, DNA binding site, intra-dimer stabilization site 1, intra-dimer stabilization site 2 and dimer-dimer stability site | $\leq 5 \text{ \AA}$ (close), $>5$ to $\leq 8 \text{ \AA}$ (near), $>8 \text{ \AA}$ (far) | 30 |

The first sequence feature included was the evolutionary conservation score calculated by ConSurf ^20^ using the ligand-bound mmpR5 crystal structure. This feature was used to assess how conserved each AA encoded by a *Rv0678* variant is based on a multiple sequence alignment of homologous proteins. The second sequence feature included was the similarity between the wild-type and a mutated protein sequence as assessed by the BLOSUM62 scoring matrix ^21^.

The one biochemical feature included was the difference in composition, polarity, and molecular volume between the AAs encoded by the wild-type and mutated *Rv0678* gene as quantified by the Grantham score ^22^.

Ten structural features were included to comprehensively assess the effect of missense *Rv0678* variants on the mmpL5 protein structure. The mCSM PPI score ^23^ was used to quantify the relative solvent accessibility by assessing how exposed a mutant residue is to solvent compared to its maximum possible exposure in an unfolded or reference state (wild-type), both in ligand-bound and DNA-bound conformation. The <u>mCSM-Stability</u> score ^23^ was used to predict how a missense variant in *Rv0678* affects the thermodynamic stability of the mmpR5 protein, either in ligand-bound or DNA-bound conformation. The mCSM-PPI2 score ^24^ was used to predict the effect of an *Rv0678* mutation on the interaction affinities between the homodimers (protein-protein) in either the ligand-bound or DNA-bound conformation. The DynaMut score ^25^ was used to assess how a missense variant in *Rv0678* affects the stability and flexibility of the mmpR5 protein in both ligand-bound and DNA-bound conformation by using normal mode analysis to predict a change in free energy. The DNA-mCSM tool ^23^ was used to assess how a *Rv0678* variant affects the affinity of the mmpR5 protein to bind to DNA in the DNA-bound conformation. To determine the shortest atomic distance to a functional site in the DNA-bound conformation, we calculated the distance between the Cα atom of a residue carrying a *Rv0678* variant and the Cα atom of each of the functional sites of interest of the *Rv0678* wild-type residue. For each variant, we then selected the shortest atomic distance (in Å) in 3D-space to any one of the functional-site positions of interest.

### Feature-phenotype association for *Rv0678* missense variants

The distribution of the 13 selected features was visualised using box plots, stratified by variants conferring a resistant or susceptible phenotype based on their majority phenotype. The association of the 13 features with variant phenotype was assessed by the Mann-Whitney U test, with p-values adjusted for false discovery rate using the Benjamini-Hochberg procedure.

### Machine learning model for feature-based phenotype prediction: development, optimization and performance assessment

We first built a binary random forest classifier to train -using the same default parameters and leave-one-out cross-validation-the published 12-feature model ^13^ on the dataset of 62 missense variants in *Rv0678* obtained through our literature review. Next, we added the new shortest atomic distance to functional sites feature to the model. To obtain the model with the minimally sufficient set of features, we removed features with the lowest mean decrease in impurity across all decision trees.

Model optimization was then performed for the reduced-feature model. Hyperparameter tuning was conducted using GridSearchCV to evaluate combinations of parameters, including the number of trees, maximum tree depth, minimum samples required for a split, minimum samples per leaf, and the number of features considered at each split. To minimise overfitting, stratified 5k-fold cross-validation was applied within GridSearchCV. This approach was preferred over leave-one-out cross-validation due to its lower variance and improved generalisation in the context of a small sample size ^26^. To reduce model complexity and remove branches with minimal contribution to predictive power, cost-complexity pruning was applied using the *ccp_alpha* parameter.

Results were aggregated across cross-validation folds to compute the area under the receiver operating characteristic curve (ROC-AUC), precision-recall area under the curve (PR-AUC), and F1 score. Overfitting of the final model was examined by measuring the accuracy gap between training and test sets through learning curves across varying training set sizes, and by using *out-of-bag* scoring to estimate how well the model would perform on test data. Interpretability of the final model was assessed using SHapley Additive explanation’s (SHAP) implemented in the *shap* Python package ^27^.

Statistical differences between the ROC-AUC values of the published 12-feature model and our final model were assessed using the DeLong test. Diagnostic performance of the final model was evaluated by deriving the point estimate and Wilson’s 95% confidence interval (95% CI) for sensitivity, specificity, negative predictive value (NPV), and positive predictive value (PPV) from the confusion matrices generated via Scikit-learn’s confusion_matrix function ^28^. This analysis was performed at both the variant level, using the unique variants retrieved from the literature review as the unit of analysis, and at the isolate level, using unique isolates as the unit of analysis.

### Sensitivity Analysis

In a sensitivity analysis, we trained a machine learning model using the same criteria as applied in the main analysis but included isolates with phenotypes determined by any phenotypic platform instead of restricting to the MGIT platform.

### External validation

The final model was applied to all isolates containing a single (SOLO) missense variant in *Rv0678* in the second version of the WHO catalogue^7^. For each unique missense variant listed in the catalogue, the majority phenotype was determined independent of the phenotypic testing platform, as the phenotypic assay used is not presented in the catalogue. Performance of the final model at the variant and isolate level was evaluated by deriving sensitivity, specificity, NPV and PPV from confusion matrices.

## RESULTS

### Isolates and *Rv0678* variants identified by the systematic review

The update of the literature search (1 January 2022 – 19 January 2026) yielded 1,585 articles (Fig. S1). After removing 482 duplicates, 1,103 articles remained. Of these, 1,029 articles were excluded based on abstract screening and 74 following full text review. The remaining 29 original research articles were added to the 49 articles identified in the prior literature review.

The 78 included articles reported 1,178 *Mtb* isolates containing 508 unique *Rv0678* variants. Of the 1,178 isolates, 1,042 isolates were excluded because they contained a synonymous or promoter region variant (n= 187), a co-occurring *atpE* variant (n=31), multiple co-occurring *Rv0678* mutations (n=143), *Rv0678* variants in positions 1–14 or 160–165 of the AA sequence (n=43), or indel or nonsense mutations in *Rv0678* (n=354). For the main analysis, an additional 276 isolates were excluded because they were tested on platforms other than MGIT or because a majority phenotype could not be assigned (n=8). The final dataset for the main analysis consisted of 136 isolates containing 62 unique missense variants in *Rv0678* of which 41 conferred a resistant and 21 a susceptible phenotype (Fig. S2).

### Identification of resistance hotspots in the 3D structure of the *mmpR5* protein

In addition to the DxR motif and Alpha-helix recognition site, three novel functional sites were identified: intra-dimer stability site 1 (residues 107-108, 151 and 154), intra-dimer stability site 2 (residues 43-46 and 103-106) and the dimer-dimer stability sites (residues 135–139) (Fig 2).

**Figure 2.**
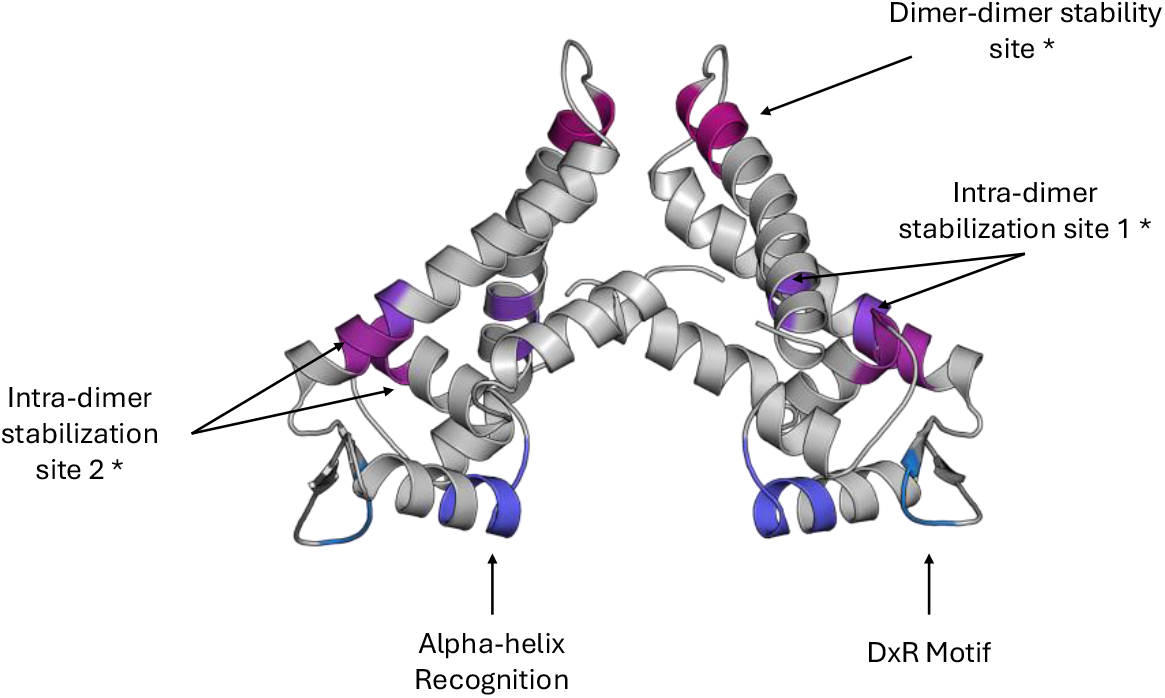
Functional sites of the mmpR5 protein in its DNA-bound conformation. *Novel sites identified by this study

Functional analysis showed that these three novel sites may play a role in ligand contact. Two sites from the intra-dimer stability site 1 lie adjacent to the aromatic residue cluster that secures the dimeric interface ^5^. Intra-dimer stability site 2 contains a position involved in salt-bridge formation, supporting its role in subunit stabilisation ^5^. The dimer–dimer stability site overlaps with the capping region of α-helix 6 and lies close to the opposing subunit in the homodimer. Given that α6 and α6′ form the antiparallel scaffold of the dimer, this positioning is consistent with a potential role in helix and protein stabilization [5, 28].

### Model development, model optimization, and model performance

The features with the strongest association with bedaquiline phenotype were evolutionary conservation score (p<0.001), chemical difference between AA (p= 0.022), shortest atomic distance to a functional site (p = 0.023), similarity between protein sequences (p = 0.033) and the protein-protein interaction in the DNA-bound conformation (p=0.237) (Fig. 3).

**Figure 3.**
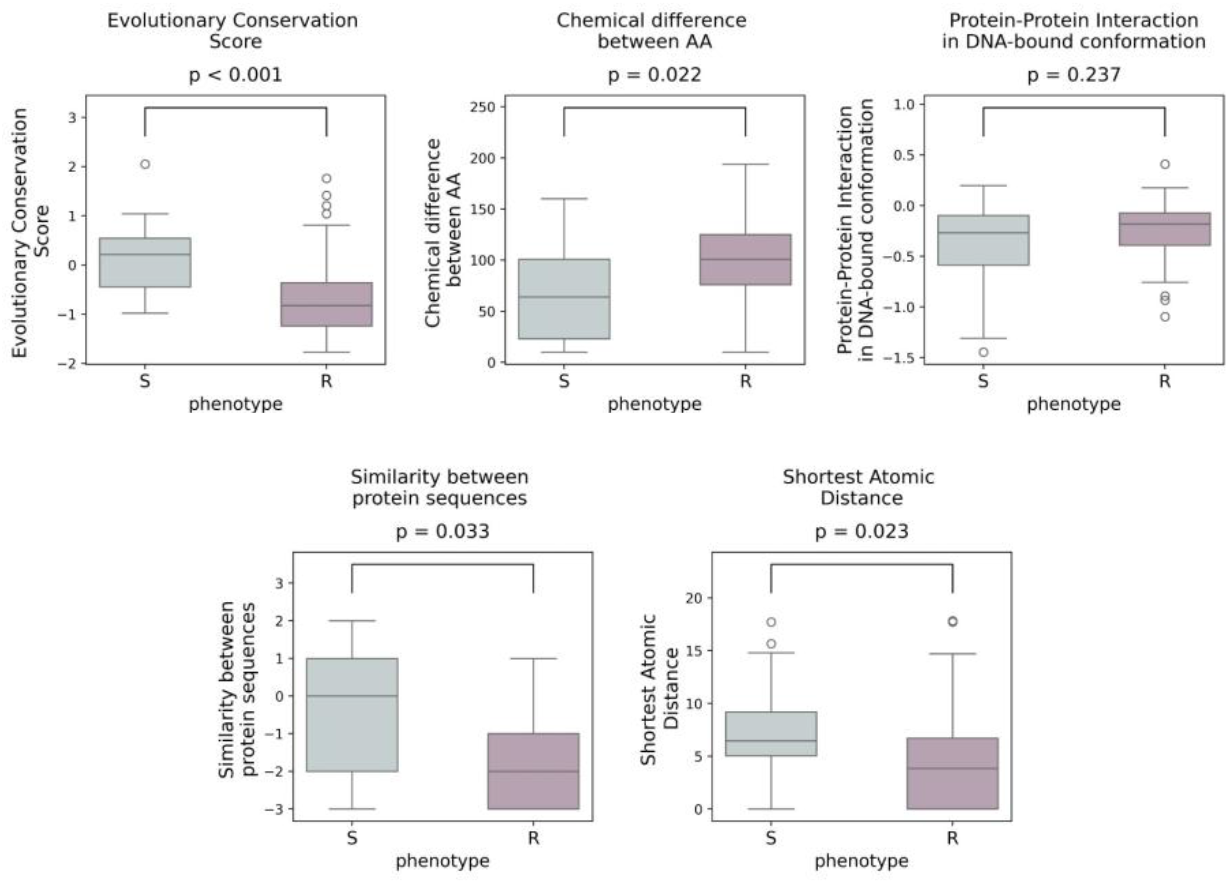
Distribution of the scores for the five most important features for 62 unique missense variants in *Rv0678* gene present in 136 clinical *Mtb* isolates, stratified by bedaquiline phenotype on MGIT.

When applied to the 62 unique *Rv0678* missense variants identified by the literature review, the published 12-feature model ^13^ achieved an AUC of 0.722 and an F1-score of 0.809. Adding the novel shortest atomic distance feature improved the model performance, increasing the AUC to 0.754 and the F1-score to 0.837. Restricting the model from 13 features to the five features with the highest mean decrease in impurity further increased the AUC to 0.801 and the F1-score to 0.880 (Table 2).

**Table 2:** Model characteristics, optimization methods and evaluation metrics of model performance.

| Model | Features | Hyper-parameter Tuning | Cross Validation method | ROC AUC | PR AUC | F1 score |
| --- | --- | --- | --- | --- | --- | --- |
| 12-feature | Evolutionary conservation score<br>Similarity between protein sequences<br>Chemical difference between AAs<br>Relative solvent accessibility - DNA-bound conformation<br>Relative solvent accessibility - ligand-bound conformation<br>Protein stability - DNA-bound conformation<br>Protein stability - ligand-bound conformation<br>Protein-protein interaction - DNA-bound conformation<br>Protein-protein interaction - ligand-bound conformation<br>Protein dynamics - DNA-bound conformation<br>Protein dynamics - ligand-bound conformation<br>Protein-DNA affinity - DNA-bound conformation | no | LOOCV | 0.722 | 0.809 | 0.822 |
| 13-feature | Same as above + shortest atomic distance to one of the 5 functional sites in DNA-bound conformation | no | LOOCV | 0.754 | 0.837 | 0.822 |
| Initial 5-feature | Evolutionary conservation score<br>Shortest atomic distance to one of five functional sites<br>Chemical difference between AAs<br>Similarity between protein sequences<br>Protein-protein interaction in DNA-bound conformation | yes | LOOCV | 0.801 | 0.880 | 0.795 |
| Final 5-feature | Same as above | yes | 5 k-fold | 0.826 | 0.887 | 0.850 |

Hyperparameter optimization yielded a configuration with 1,000 decision trees, a maximum tree depth of 5, the square root of the total features used at each split, minimum samples required to split a node set to two, and minimum samples required to form a leaf set to ten. The optimized five-parameter model demonstrated overfitting, with a gap between training (mean accuracy = 0.796) and test (mean accuracy = 0.923) learning curves (Fig. S3). 5 K-fold cross-validation successfully minimised overfitting, reducing the gap between training and testing learning curves from 0.127 to 0.017 (Fig. S4). After overfitting corrections, the out-of-bag error curve stabilized after approximately 100 trees (Fig. S5).

The final 5-feature model achieved an F1-score of 0.826 (Fig. S6) and increased the AUC from 0.722 for the published 12-feature model to 0.826 (p =0.00026) (Fig. S7). SHAP analysis of the final model showed that evolutionary conservation and shortest atomic distance were the strongest contributing features to resistance prediction, with low scores associated with positive SHAP values, reflecting bedaquiline resistance (Fig. 4, Fig S8).

**Figure 4.**
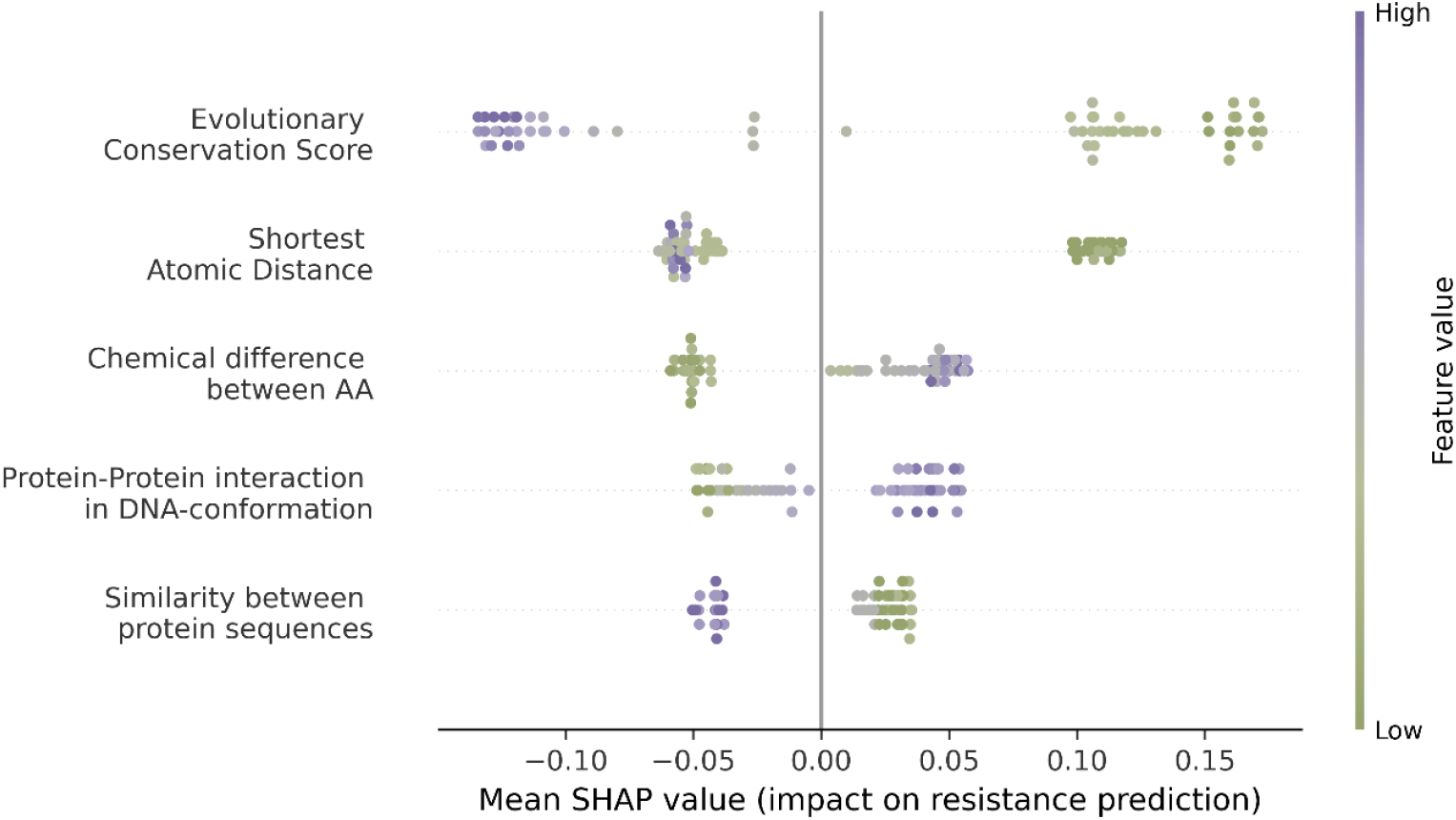
SHAP beeswarm plot visualizing the contribution of the 5 features included in the final model on the bedaquiline resistant phenotype prediction. Each dot represents a missense variant in *Rv0678*, the position on the x-axis represents the SHAP value (i.e. impact on model prediction), the color represents the feature’s value (green = low, purple = high). SHAP values with a high absolute positive value indicate a substantial contribution to the model’s resistance prediction.

### Sensitivity, specificity PPV and NPV of the final prediction model

At the variant level (including 62 unique missense variants), the final five-feature model showed a sensitivity of 87.8% (95% CI, 74.5–94.7), specificity of 81.0% (95% CI, 60.0– 92.3), PPV of 90.0% (95% CI, 76.9–96.0), and NPV of 77.3% (95% CI, 56.6–89.9). At the isolate level, the performance of the final five-feature model was assessed using the 136 isolates, with a sensitivity of 87.1% (95% CI, 78.3–92.6), specificity of 88.2% (95% CI, 76.6–94.5), PPV of 92.5% (95% CI, 84.6–96.5), and NPV of 80.4% (95% CI, 68.2– 88.7) (Table S1).

### Sensitivity Analysis

For the sensitivity analysis, 141 unique missense variants in *Rv0678* and 303 isolates with phenotypic information obtained by any DST platform were included. Of the 141 variants, 73 had a resistant and 68 a susceptible bedaquiline phenotype (Fig. S9). Because the model trained on this dataset had a poor discrimination between resistant and susceptible phenotypes (AUC 0.649) (Fig. S10), we did not perform further assessments.

### External model validation

In the second version of the WHO catalogue, 159 SOLO *Rv0678* missense variants are listed in 362 isolates. Of the 159 variants, 26 were excluded from the analysis because their location at AA positions was not represented in the mmpR5 crystal structure (n = 10) or because a majority phenotype could not be assigned (n = 16), resulting in 133 unique variants occurring in 342 isolates. For these 133 unique *Rv0678* missense variants, model sensitivity was 71.6% (95% CI, 60.5–80.6), specificity 47.5% (95% CI, 35.3– 60.0), PPV 63.1% (95% CI, 52.4–72.6), and NPV 57.1% (95% CI, 43.3–70.0). At the isolate level, sensitivity was 66.7% (95% CI, 59.5–73.1), specificity 59.9% (95% CI, 52.2–67.1), PPV 64.9% (95% CI, 57.7–71.4), and NPV 61.8% (95% CI, 54.0–69.0) (Table S2).

## DISCUSSION

Early detection of resistance to bedaquiline is critical to achieve good treatment outcomes in patients suffering from RR-TB. We show that a novel *in-silico* feature-based model built using machine learning methods achieved a high discriminator power (AUC 0.826) and could predict the bedaquiline phenotype of a missense variant in *Rv0678* with 87.1% sensitivity and 88.2% specificity. Evolutionary conservation score and shortest atomic distance to functional sites were the most important features for predicting the bedaquiline phenotype.

Three in silico models have been published previously. Compared to a 12-feature model for *Rv0678* missense variants, our model performed significantly better (AUC 0.826 vs 0. 722), likely due to the addition of a new feature of distance to a functional site and use of feature reduction methods, which improved model robustness through higher dimensionality (5 vs 12 features) relative to the sample size ^13^. The model for *atpE* variants ^12^ achieved even higher accuracy (AUC 0.93), likely because the deleterious effects of *atpE* mutations result in very high MICs and thus more pronounced differences between wild-type and mutant isolates. Similar to our model, informative features for the *atpE* model were proximity to a critical functional region of the protein and predicted effect of the mutation on the AA sequence, albeit measured by SNAP2 to predict the functional effects of variants instead of directly assessing evolutionary conservation. There were also important differences. Protein stability in ligand conformation (mCSM-Lig) was highly predictive in the *atpE* model, but showed limited discriminatory power in our *Rv0678* model, likely because *atpE* acts directly on the bedaquiline binding site, whereas *Rv0678* alters the regulatory function of the mmpS5/mmpL5 efflux pump. Compared with our model, the Convolutional Neural Network showed lower sensitivity for predicting bedaquiline resistance across resistance genes (46.2%). Both approaches leverage sequence-based information; however, our model identified an explicit biochemical descriptor (Grantham distance) as an important contributor, whereas the CNN’s composite biochemical features based on molecular weight, isoelectric point, and hydrophobicity contributed minimally.

Results of our study need to be interpreted in light of several limitations. First, while multiple co-occurring variants occur in *Rv0678*, our model can only predict the phenotypic effect of single missense variants ^15^. Second, our model cannot predict the phenotype of nonsense or loss-of-function (LoF). variants in *Rv0678*. The prediction of the phenotype of LoF is, however, of less clinical importance as the WHO grading rules classify all LoF variants as conferring bedaquiline resistance ^7^. Third, while the identification of novel functional sites related to dimerization stability expands our understanding of the mmpR5 protein, for which analyses have mostly focused on the DNA-binding regions, the new functional sites should be interpreted with caution, as these have not been validated through *in vitro* studies. Finally, the model performance for variants and isolates listed in the WHO catalogue was substantially lower. We hypothesize this is due to the measurement error introduced when using multiple pDST platforms, including platforms for which there is no standardized critical concentration threshold, such as colorimetric assays or solid media-based methods. This hypothesis is supported by our sensitivity analysis, which showed that including data from any pDST platform (rather than restricting to only the MGIT platform) resulted in poor performance (AUC = 0.649) of our final model. Similarly, the prior 12-feature *Rv0678* model also reported reduced performance when using multiple pDST platforms ^13^. The low performance of the Convolutional Neural Network may also have been driven by the inclusion of multi-platform WHO catalogue data ^14^

In conclusion, the high discriminatory power and accuracy of an *in-silico* feature-based model highlight the power of using information on the molecular and biological consequences of genomic variants for the prediction of the bedaquiline resistance phenotype. Integration of this model into variant interpretation software could enhance the clinical interpretation of missense variants in *Rv0678* and thus improve the ability of sequencing methods to guide treatment in a clinical context. Future research should update the model as more paired genotype-phenotype data on bedaquiline becomes available, as machine learning methods improve, and as new tools for protein structure prediction emerge.

## AUTHOR CONTRIBUTIONS STATEMENT

W.Q.R. updated the literature review, developed and optimized the model, performed data analysis and wrote the manuscript. M.d.D.F. performed the mmpR5 analysis and feature quantification. V.R. performed graphical visualization and mmpR5 analysis, and edited the manuscript. E.R. developed the baseline model, conducted the initial literature review and supported model development. M.S. contributed expert input and guidance for model optimization. A.V.R. supervised the study and contributed to writing and editing of the manuscript.

## Notes

### Competing Interest Statement

The authors have declared no competing interest.

