## Supplementary Material for "Feature-based in-silico model to predict the *Mycobacterium tuberculosis* bedaquiline phenotype associated with Rv0678 variants"

### SUPPLEMENTARY MATERIALS

#### List of supplementary figures


| | |
|---|---|
| Fig S1 | Flow diagram of the systematic review. |
| Fig S2 | Selection of isolates and unique variants for the main analysis (MGIT platform). |
| Fig S3 | Learning curves of the initial 5-feature model. |
| Fig S4 | Learning curves of the final 5-feature model. |
| Fig S5 | The out-of-bag (OOB) error curve of the final 5-feature model. |
| Fig S6 | Performance of the final 5-feature model. |
| Fig S7 | ROC curves comparison between the 12-feature baseline model and the final 5-feature model. |
| Fig S8 | Mean absolute SHAP values for the five features included in the final 5-feature model. |
| Fig S9 | Selection of isolates and unique variants for the sensitivity analysis. |
| Fig S10 | Sensitivity analysis in all pDST dataset. |


#### List of supplementary tables


| | |
|---|---|
| Table S1 | Model predictions for variants identified by the systematic literature review. |
| Table S2 | Model performance for variants listed in the WHO Mutation Catalogue |

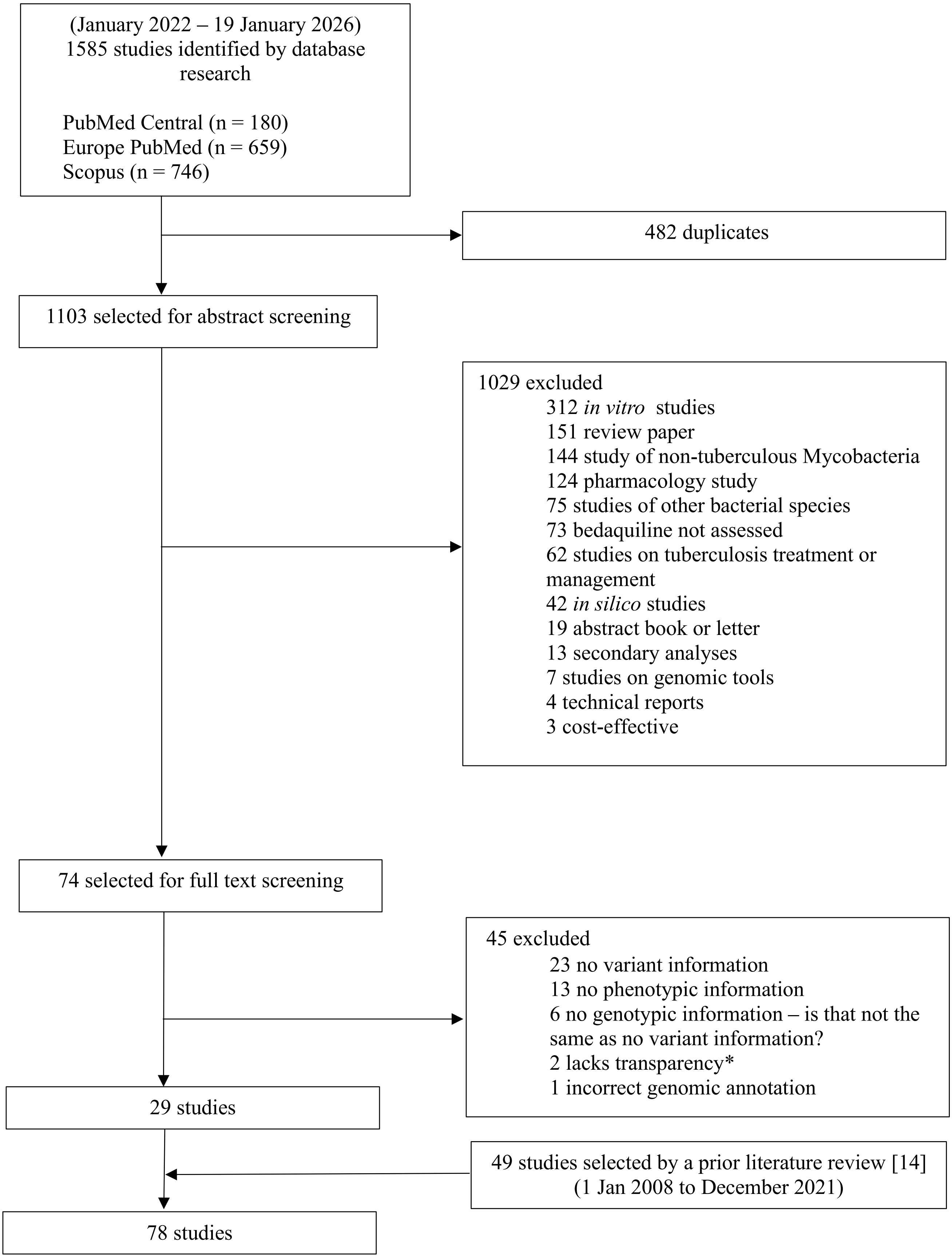


Figure S1: Flow diagram of the systematic literature review. *Unclear methodology section and/or references to unavailable data.

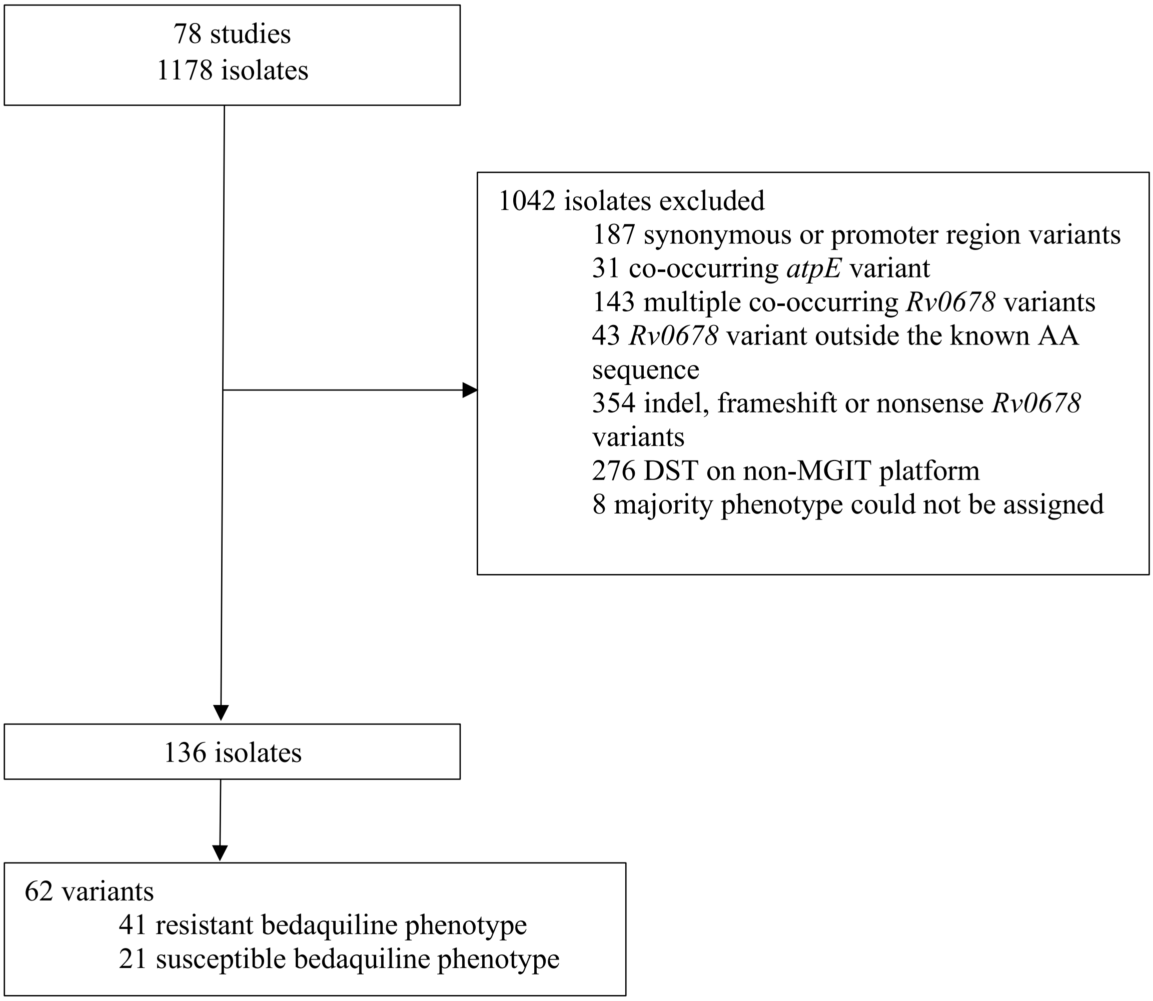


Figure S2: Selection of isolates and unique variants for the main analysis, restricted to phenotypic data obtained by the MGIT platform

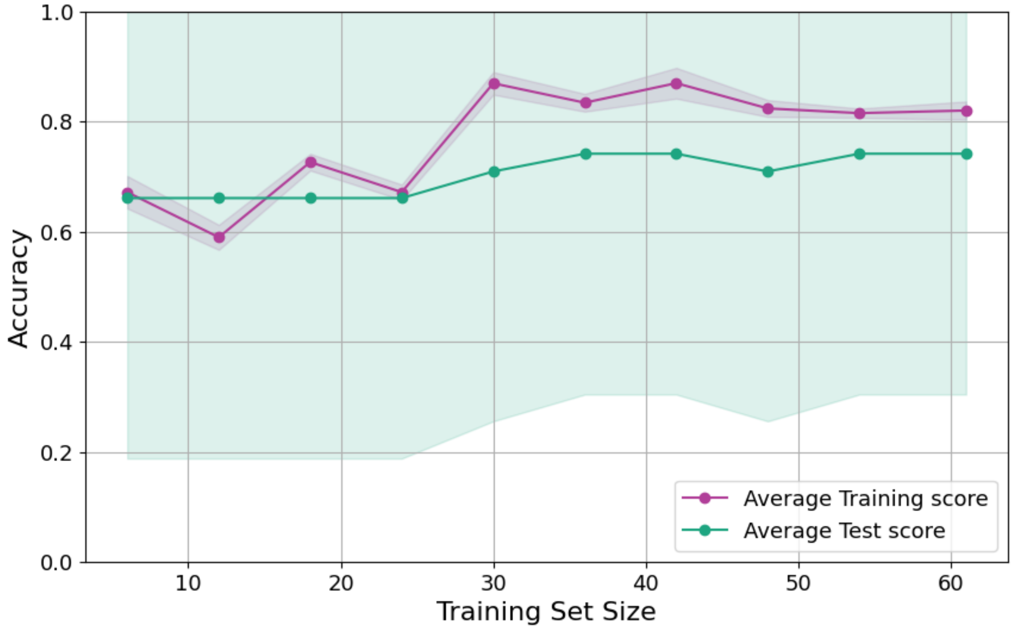


Figure S3: Learning curves of the initial 5-feature model using LOOCV and Grid Search hyperparameter tuning. Average train accuracy = 0.796 and average test accuracy = 0.923

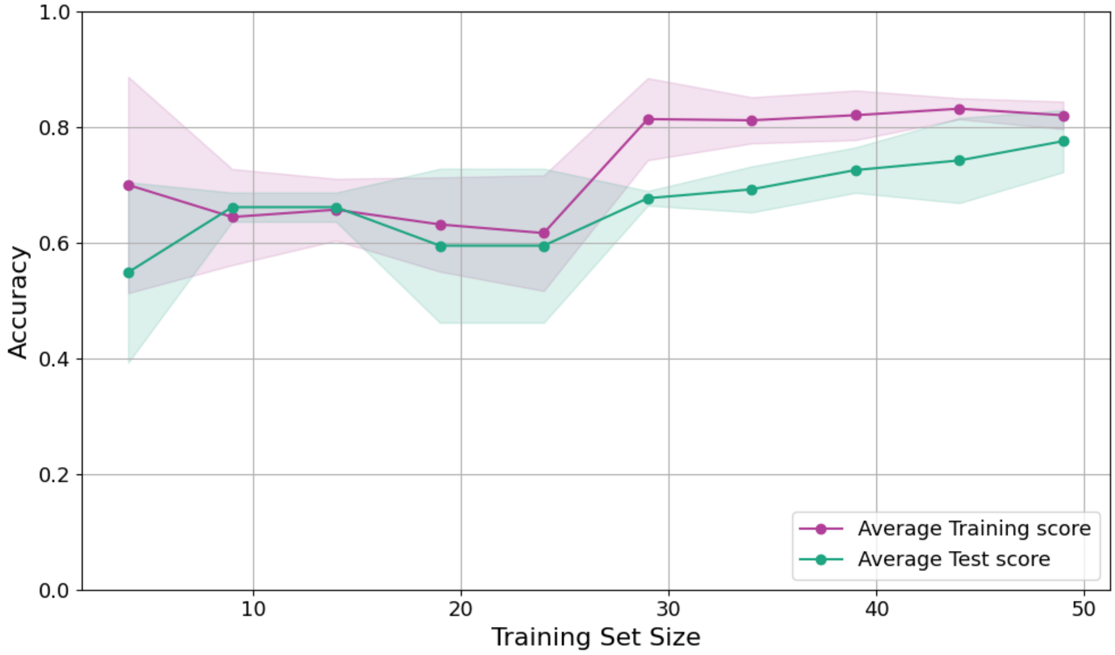


Figure S4: Learning curve of the final 5-feature. Average train accuracy = 0.823 and average test accuracy = 0.806

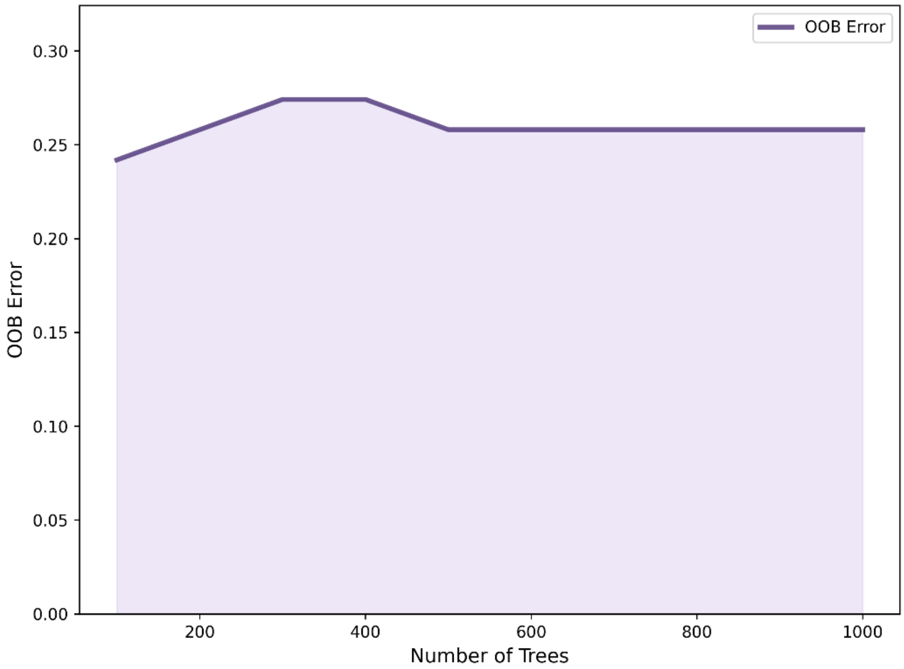


Figure S5: The out-of-bag (OOB) error curve of the final 5-feature model using K-fold cross-validation.

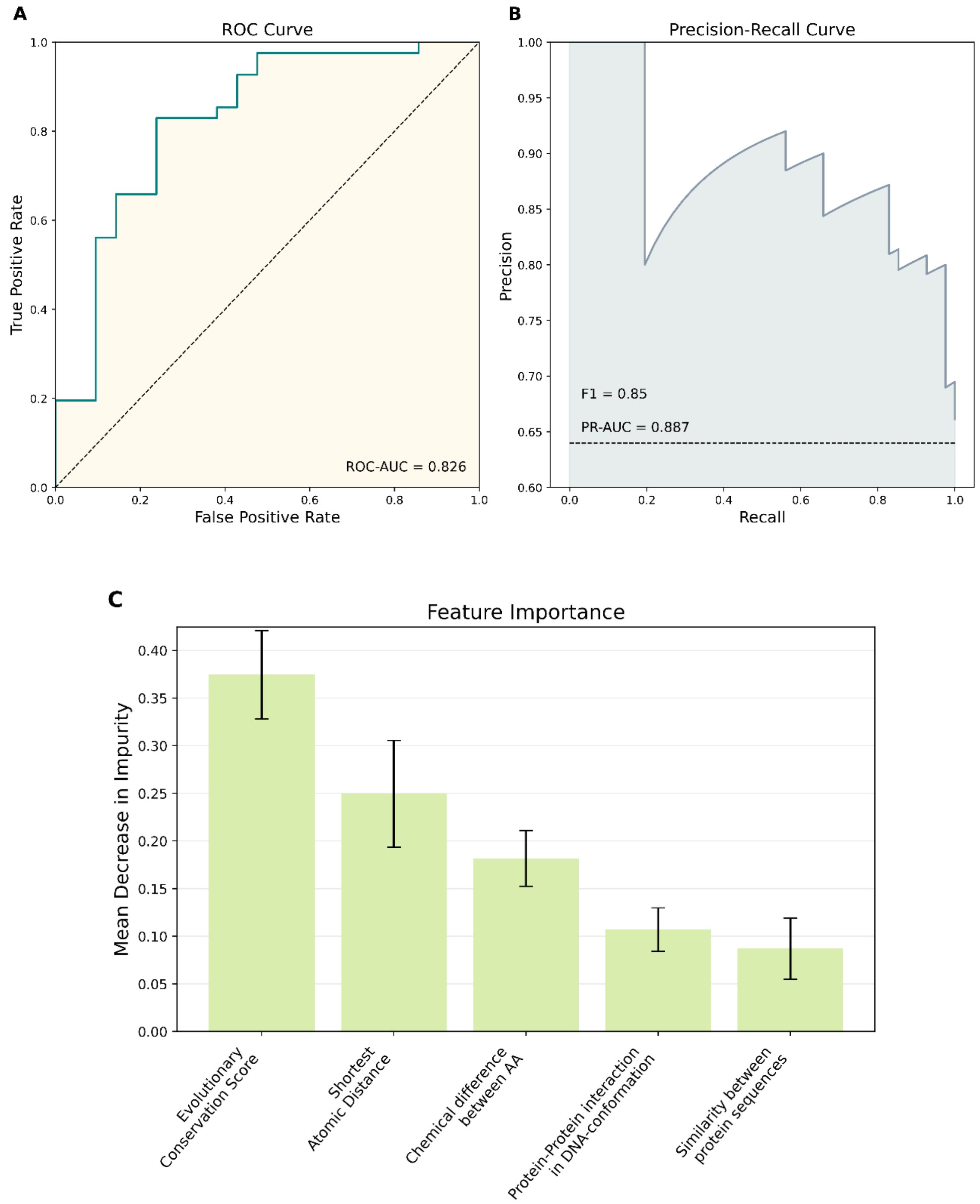


Figure S6: Performance of the final 5-feature model. (A) Receiver operating characteristic (ROC) curve illustrating the model's ability to discriminate between resistant and susceptible phenotypes; the diagonal dashed line represents random classification. (B) Precision–recall curve showing performance under class imbalance. (C) Feature importance ranked by mean decrease in impurity.

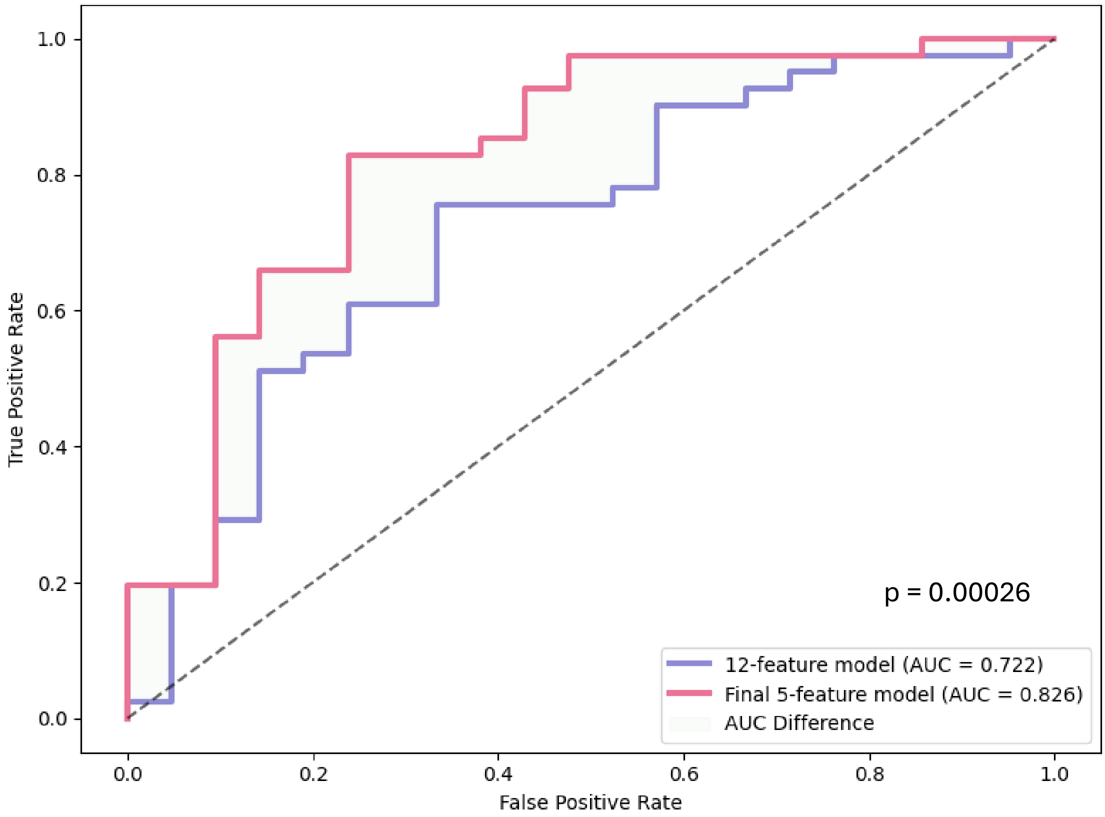


Figure S7: ROC curves comparison between the 12-feature baseline model and the final 5-feature model. P value obtained by DeLong test

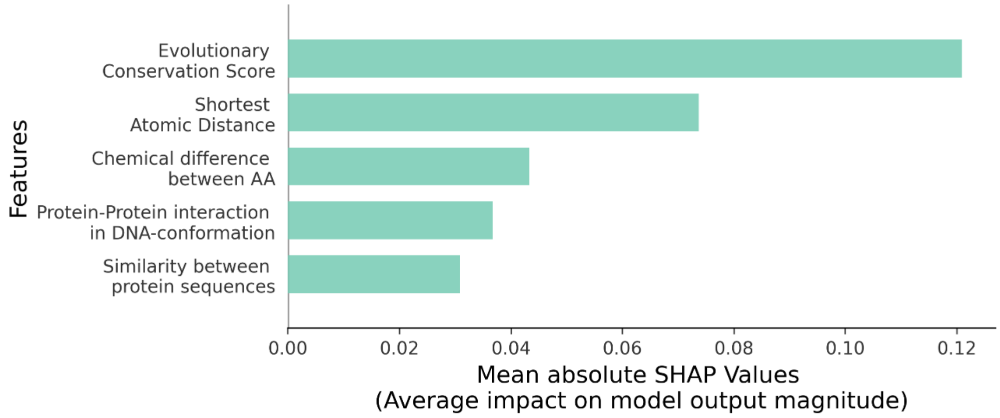


Figure S8: Mean absolute SHAP values for the five features included in the final 5-feature model. The SHAP value represents the average contribution to the model's prediction of bedaquiline resistance.

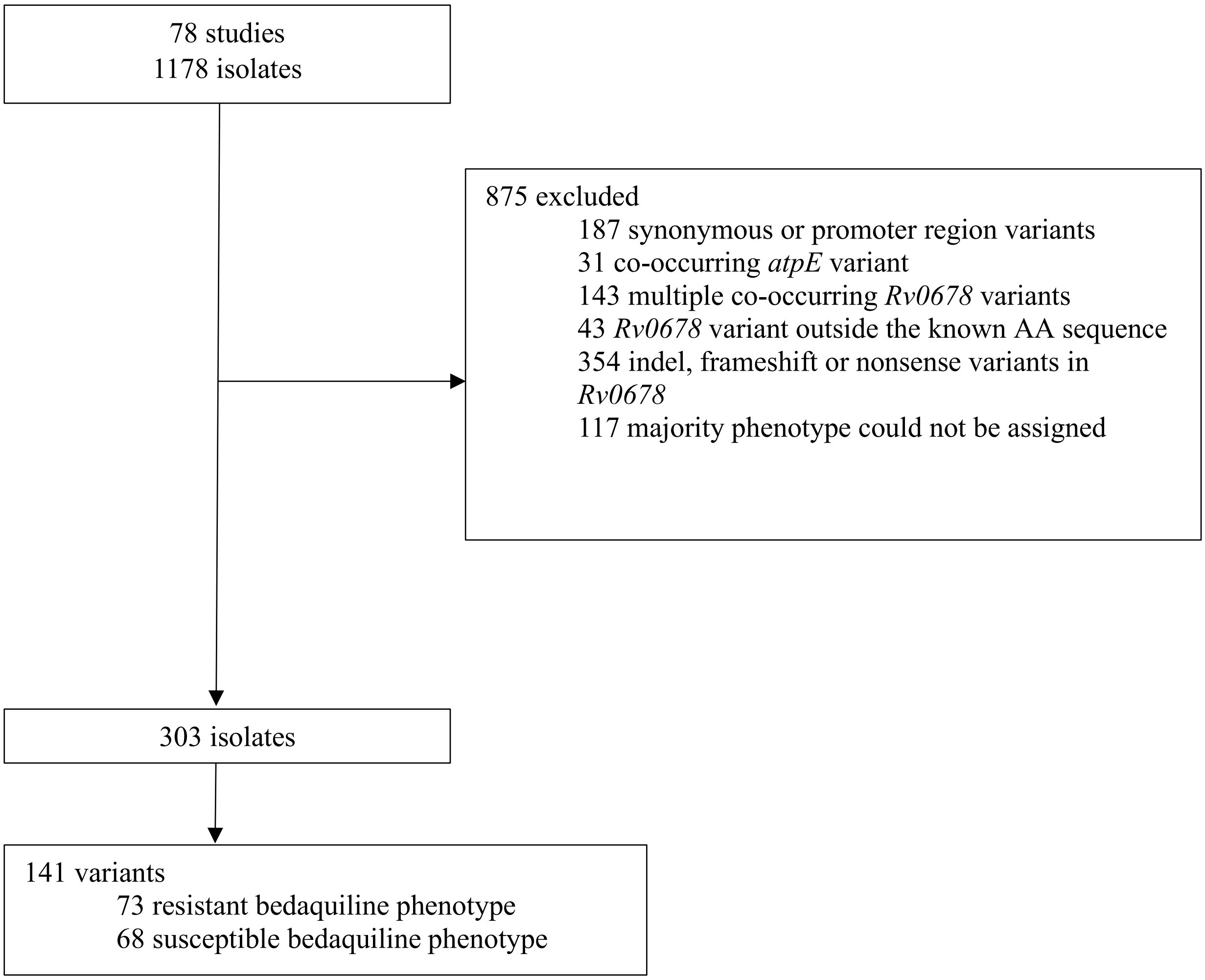


Figure S9: Selection of isolates and unique variants for the sensitivity analysis using any phenotypic DST platform

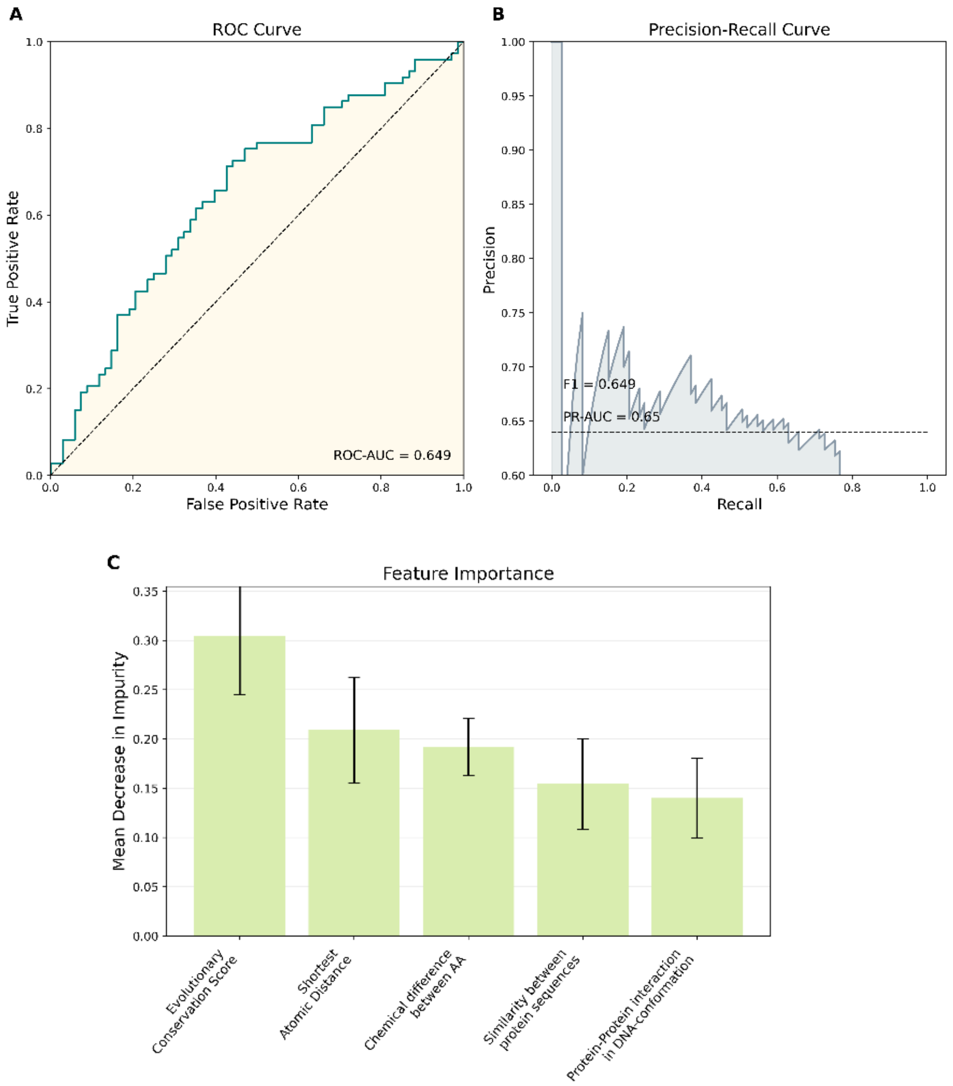


Figure S10: Performance of the final five-feature model trained on the sensitivity dataset (all pDST platforms). A) ROC curve, B) Precision-Recall Curve, and C) feature importance.

Table S1: Model predictions for 62 unique *Rv0678* variants and 136 clinical isolates identified by the systematic literature review.

| ***Phenotype*** | Majority Resistant phenotype on MGIT | Majority Susceptible phenotype on MGIT |
|---|---|---|
| 62 unique *Rv0678* missense variants | | |
| ***Model prediction*** | | |
| Resistant | 36 | 4 |
| Susceptible | 5 | 18 |
| 136 isolates containing a single missense variant in *Rv0678* | | |
| ***Model prediction*** | | |
| Resistant | 74 | 6 |
| Susceptible | 11 | 45 |

Table S2: Model performance for 133 unique *Rv0678* variants and 342 clinical isolates listed in the WHO Mutation Catalogue (2nd Edition).

| ***Phenotype*** | Majority phenotype Resistant | Majority phenotype susceptible |
|---|---|---|
| 133 unique missense variants in *Rv0678* | | |
| ***Model prediction*** | | |
| Resistant | 53 | 31 |
| Susceptible | 21 | 28 |
| 342 isolates containing a single missense variant in *Rv0678* | | |
| ***Model prediction*** | | |
| Resistant | 120 | 65 |
| Susceptible | 60 | 97 |
